# Behavioral response of bed bugs (Hemiptera: Cimicidae) to wet surfaces

**DOI:** 10.1101/2025.07.31.667989

**Authors:** Jorge Bustamante, Dong-Hwan Choe

## Abstract

The behavioral response of bed bugs (*Cimex lectularius*) to wet surfaces was observed using motion-tracking software in a circular arena with a filter paper floor. Half of the arena floor was treated with varying amounts of deionized water, resulting in four treatment conditions: no water (control), low, medium, or high application rates. The other half was left untreated. For both sexes (adult) and all nymphal stages, the bed bugs spent significantly less time and walked significantly shorter distances on the wet surface. This result was amplified with increased water application rates. Some bed bugs made turns as they approached the wet zone. Bed bugs increased their speed by 38% when fleeing the wet zone relative to their approaching speed. On average, bed bugs made turns when they were 0.58 cm away from the wet zone. Furthermore, nymphs made turns when they were farther away from the wet zone relative to adults (by 60% on average). These findings suggest that overapplication of liquid pesticides may cause bed bugs to avoid treated areas, highlighting the need for precise application rates to enhance control efficacy. Additionally, this avoidance behavior likely reflects an adaptive strategy to reduce the risk of fungal infection or drowning, consistent with the evolutionary history of bed bugs as cave-dwelling insects.

## Introduction

The common bed bug *Cimex lectularius* (L.) and the tropical bed bug *C. hemipterus* (F.) are the most widespread bed bug pests for humans, and they are obligate blood feeders at all life stages (Usinger 1966; Doggett et al. 2018; Doggett and Lee 2023). A global resurgence of bed bugs has occurred for the past twenty years (Doggett et al. 2018). Many attributes, including their small size (4 mm body length in adulthood) and behavior, contribute to the difficulty of detecting and monitoring bed bugs (Usinger 1966; Vaidyanathan and Feldlaufer 2013; Wang et al. 2018; Crawley and Borden 2021). The evolution of resistance to chemical pesticides directly contributes to the complexity of bed bug control, as many field strains of bed bugs are now resistant to common, widely available insecticides (Dang et al. 2017). Bed bugs also exhibit behavioral responses to substrate properties, such as texture and chemical treatments, which can influence their resting preferences and movement patterns (Hottel et al. 2015). Therefore, gaining a deeper understanding of their behavior is essential for improving monitoring, detection, and control methods.

All extant Cimicidae likely evolved from ancestral species associated with hosts nesting in elevated habitats (Usinger 1966; Roth et al. 2019). Many species inhabit caves and aggregate in rooftop crevices, avoiding cave floors where water can accumulate (Usinger 1966). An extant example is the African bat bug (*Afrocimex constrictus*), which inhabits caves with 60-80% relative humidity (RH) and aggregates away from cave floors (Reinhardt et al. 2008). Similarly, cimicids that feed on birds also avoid areas where water can accumulate (Usinger 1966). These observations suggest that bed bugs may avoid water accumulation, possibly through the ability to detect moisture or associated environmental cues.

The ability of an organism to detect humidity or moisture in the environment occurs in many insects and is mediated by specialized hygroreceptors (Altner and Loftus 1985). This includes but is not limited to insects in Blattodea (Roth and Willis 1952; Yokohari 1978), Coleoptera (Roth and Willis 1951; Harbach and Larsen 1977), Diptera (Roth and Willis 1952; Kellogg 1970), Hymenoptera (Yokohari et al. 1982; Steidle and Reinhard 2003), and Lepidoptera (Steinbrecht and Müller 1991; Rowley and Hanson 2007). Other non-insect arthropods also demonstrate hygroreception (Gunn 1937; Ehn and Tichy 1994; Krober and Guerin 1999; Hayward et al. 2000; Kröber and Guerin 2000). In bed bugs, hygroreceptors are located on the antennae (Steinbrecht and Müller 1976).

Bed bug nymphs often die after molting at 0-20% RH (Kemper 1936), and adult females exhibit reduced longevity at ≤ 33% RH (Benoit et al. 2009). Similarly, when RH approaches 100%, bed bug survival is significantly reduced (Omori 1941), likely due to lethal fungal infection (Kemper 1936).

Other hematophagous arthropods also demonstrate hygroreception. Bugs in the family Reduviidae detect and respond to water vapor during host detection (Guarneri et al. 2002; Barrozo et al. 2003; Indacochea et al. 2017). Because heat affects the capacity of air to hold moisture, *Triatoma infestans* use humidity as a short-range directional cue for warmth, presumably emitted by vertebrate hosts (Barrozo et al. 2003). *Triatoma* nymphs prefer 30% RH and avoid 90% RH (Indacochea et al. 2017), indicating a broad detection range. At < 9.3% and ≥ 99.9% RH, egg hatching and molting by first instar *T. brasiliensis* are reduced, and nymphs avoid > 35.9% RH immediately after feeding (Guarneri et al. 2002).

Ixodid ticks require water vapor to replenish their water deficit but avoid contact with water to prevent ingesting pathogens (Kahl and Alidousti 1997; Kröber and Guerin 2000). Kröber and Guerin (2000) observed larvae of *Amblyomma variegatum, Boophilus microplus*, and *Ixodes ricinus* (including nymphs) turn away from a wet membrane upon contact, walk along the border between wet and dry membranes, and eventually settle on the wet surface.

After observing adult male *C. lectularius* approaching a wet surface and turning away approximately 1 cm before making contact, Rivnay (1932) speculated that bed bugs characteristically avoid wet surfaces. If bed bugs are placed on a wet surface, they walk in a “stilted” manner with legs extended and the abdomen lifted (Hase 1917). Despite these observations, the ability of bed bugs to detect water and their behavioral response to wet surfaces has not been experimentally verified. This study investigates the orientation mechanisms of bed bugs in response to wet surfaces, focusing on their ability to detect and avoid moisture through negative hydrotaxis. We characterized the behavioral responses of *C. lectularius* when approaching or in contact with surfaces holding varying amounts of water.

## Materials and Methods

### Bed bugs

Earl Strain bed bugs (*C. lectularius*) were originally obtained as adult males and females from Sierra Research Laboratories (Modesto, CA, USA) in 2014. All bed bugs were fed on defibrinated rabbit blood (HemoStat Laboratories, Dixon, CA, USA) using an artificial feeding system. Colonies were maintained at 24–26°C and 15–30% RH, with a photoperiod of 12:12 (L:D) hr in screened vials (4.5 cm diameter × 9 cm height) lined with corrugated filter paper for harborage. The bed bugs were used for behavioral assays within 14 d from the most recent blood meal (mean ± SE: 5.4 ± 0.35 days, *n* = 108 bed bugs).

### Experimental arena

The experimental arena was built by placing an acrylic ring wall (10.8 cm outer diameter, 10.2 cm inner diameter, 3 cm high) on two halves of a 12.5 cm diameter filter paper disc (Whatman Qualitative filter paper, Cat. No. 1001 125, Sigma-Aldrich, Merck KGaA, Darmstadt, Germany). An inverted tissue culture dish lid (Falcon® 150 mm Cell Culture Dish with 20 mm molded grid, TC-treated polystyrene, Corning Inc. Life Sciences, Durham, NC, USA) served as the arena floor (Figure 1A). The weight of the ring wall kept the pieces of filter paper flat and flush with the arena floor. The inner and outer surfaces of the ring wall were coated with Fluon (Teflon PTFE DISP 30 – 500 mL, Fuel Cell Earth LLC, Stoneham, MA, USA) to prevent bed bug escape. The experimental arena room temperature ranged from 24–26°C and 15–30% RH.

**Fig. 1.**
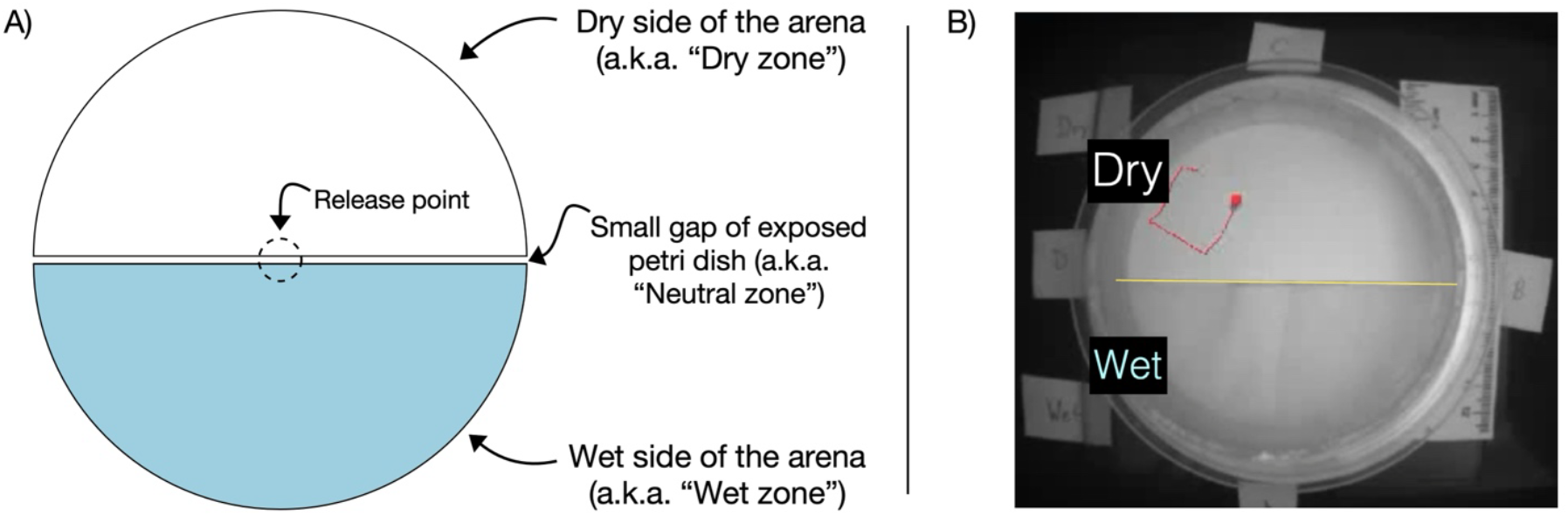
Design of arena for bed bug motion tracking. **A**. Schematic of the arena divided into wet and dry halves, with a small neutral zone gap of exposed petri dish between the two halves and a central release point. **B**. Still image of the arena as recorded by EthoVision. The red line indicates 10 s of a bed bug path, starting at the top, turning away from the wet zone, and ending at the red dot.

A small 1-2 mm wide gap (the “neutral” zone) separated the 61.35 cm^2^ wet and dry zones (Figure 1A). De-ionized water was applied to the wet zone at 0.008, 0.016, and 0.024 mL/cm^2^ (low, medium, and high application rates, respectively) (Figure 1A) and allowed to permeate the entire surface for 2-10 s before assays began. The dry zone was left untreated. In the control treatment, both zones were dry. The positions of wet and dry zones were alternated between trials at each application rate.

### Motion tracking

The movement of bed bugs was recorded with a near-infrared camera (Basler, acA1300-60gmNIR, Ahrensburg, Schleswig-Holstein, Germany) in a dark room illuminated with an infrared illuminator (Axton AT-11S, 850 nm, 150^°^, North Salt Lake, UT, USA) without any visible light sources. Bed bugs (nine each of adult males, adult females, and nymphs per treatment) were placed individually in the center of the arena, confined within an upright glass tube (1 cm internal diameter x 1 cm high) to prevent pre-trial escape. Recording for 10 min began immediately after removing the tube (Figure 1B). The primary author was present in the room to induce the movement of bugs exposed to exhaled CO_2_. Video recordings were analyzed by dynamic subtraction with EthoVision XT version 11.5 (Noldus Information Technology, Wageningen, The Netherlands), with contrast manually adjusted to distinguish a bed bug from the filter paper surface. EthoVision recorded the Cartesian *x and y* positions of bed bugs to provide kinematic metrics such as the total distance traveled, cumulative duration spent, and the mean walking speed in wet, dry, and neutral zones.

### Turning behavior analysis

Some bed bugs in the dry zone turned away from the wet zone after approaching it. An algorithm (GitHub: https://github.com/JorgeBJr) was developed to analyze the continuous path walked as a bed bug approached the wet zone. The path was defined by continuous *x* and *y* Cartesian points within four adult bed bug body lengths (4 × 0.4 cm) of the wet zone border, based on the raw data coordinates. Approach and flee sections of this path were determined based on the following requirements: exactly one minimum distance point to the wet zone and at least 100 frames (the equivalent of 1.46 s) of data on both sides of the frame associated with the minimum distance point. Because each path required a minimum of 201 contiguous frames, the total minimum time of each path must be at least 2.93 s.

The mean speeds of the bed bugs as they approached (*v*_*approach*_) and walked away from (*v*_*flee*_) the wet zone after turning within the minimum distance were calculated from 100 frames each. The speed difference (*v*_*flee*_ *-v*_*approach*_) was compared with zero (null hypothesis: no difference) to test whether bed bugs consistently fled faster than they approached. To assess whether this speed change varied with application rate, speed differences were compared across application rates using a Kruskal-Wallis test.

### Statistical analysis

Statistical analyses (*α* = 0.05) were performed using RStudio (Version 2023.06.0+421). Because Shapiro-Wilk tests indicated non-normal distributions, responses between wet and dry zones for data pooled for nymphs and both adult sexes were analyzed separately for low, medium, and high water application rates using the Wilcoxon signed-rank test. Data from all application rates were pooled to determine if the *v*_*flee*_ *-v*_*approach*_ differed from zero (if the null hypothesis were true) using a one-sample Wilcoxon signed-rank test. Comparisons between application rates for speed difference, minimum distance to the wet zone, slopes of the minimum distance to the wet zone, and number of approaches to the wet zone were conducted using Kruskal-Wallis tests followed by Dunn’s tests for multiple comparisons with a Bonferroni correction when necessary for the minimum distance to the wet zone. The Wilcoxon rank-sum test was employed to compare the minimum distance to the wet zone between adults and nymphs.

For metrics such as speed difference (*v*_*flee*_ *-v*_*approach*_) and minimum distance to the wet zone, turns were pooled across individuals within each application rate. Non-parametric tests (e.g., Wilcoxon signed-rank test, Kruskal-Wallis test) were used to account for potential intra-individual variability, as these are robust to dependencies within individuals. Additionally, a supplementary mixed-effects model was performed to confirm that the results were consistent when explicitly accounting for the nested structure of the data (turns nested within individual bugs).

For comparisons between complementary data (e.g., time spent on wet vs. dry sides of the arena), we used paired non-parametric tests (Wilcoxon signed-rank test) to account for the paired nature of the data. Each bed bug was treated as an independent data point, and the time spent on the wet side was directly compared to the time spent on the dry side for the same individual. This approach ensures that the assumption of independence is not violated and that differences are not overestimated.

## Results

### Bed bugs avoid the wet zone

Bed bugs walked significantly shorter distances on the wet than dry zones for all three application rates (Wilcoxon signed-rank exact test, low: *V* = 356, *P* < 0.0001, medium: *V* = 378, *P* < 0.0001, high: *V* = 378, *P* < 0.0001, control *V* = 230, *P* = 0.34). The median (interquartile range, IQR; 25th–75th percentiles) distances traveled for wet and dry zones, respectively, were 2.47 (IQR: 0-27.7) *vs*. 120.9 (IQR: 79.6-187.0) cm (low), 0 (IQR: 0-0) *vs*. 131.5 (IQR: 76.2-188.6) cm (medium), 0 (IQR: 0-0.8) *vs*. 122.5 (IQR: 79.6-204.4) cm (high), 111.5 (IQR: 37.4-142.3) *vs*. 90.9 (IQR: 55.0-129.8) (control) (Figure 2A).

**Fig. 2.**
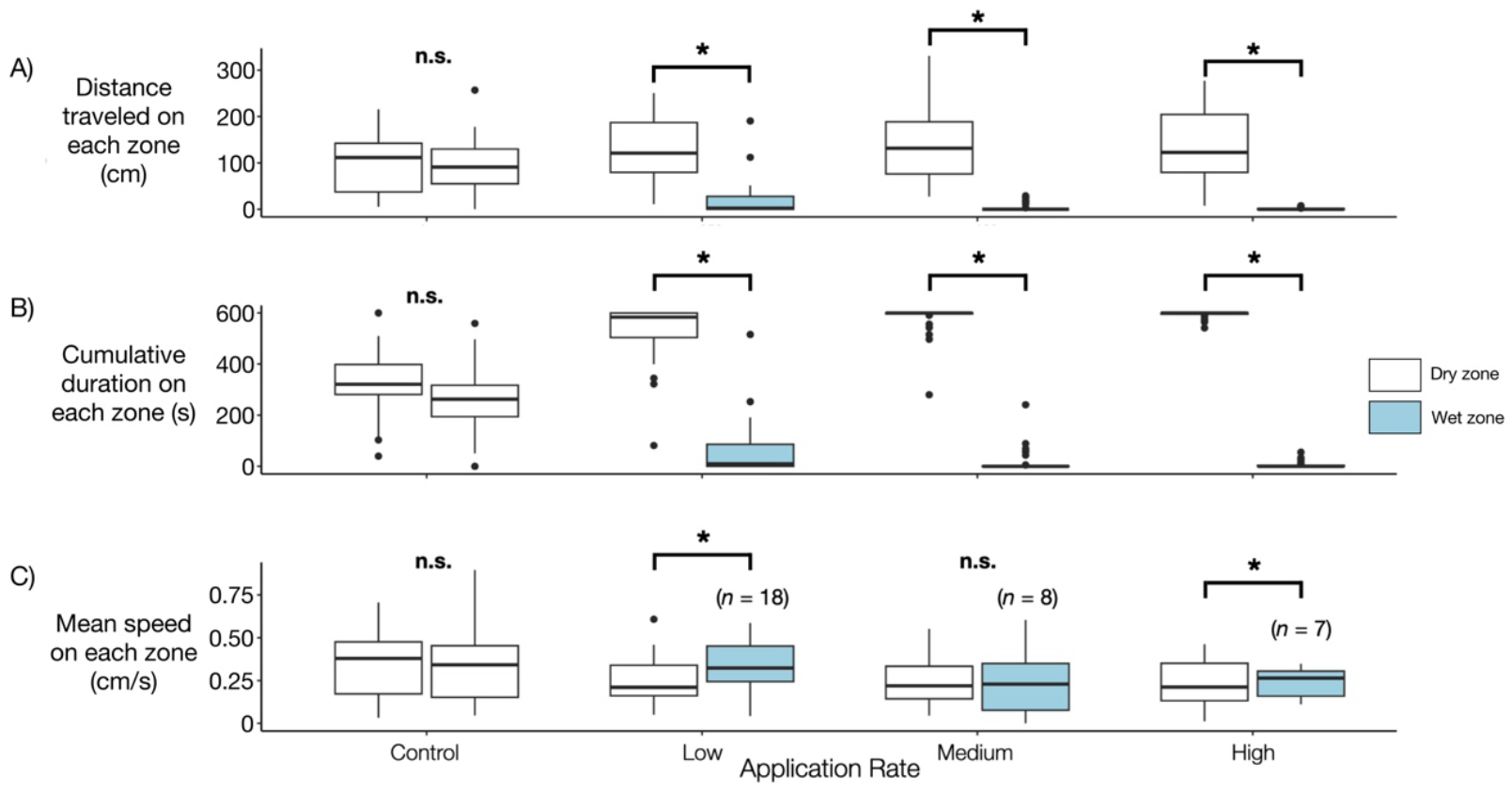
Kinematics of bed bugs (data pooled for nine each of nymphs and male and female adults) walking on different halves of the arena. **A**. Distance traveled on each side of the arena. **B**. Total time spent on each side of the arena. **C**. Mean speed on each zone. Comparisons with fewer than 27 bed bugs were due to bed bugs not contacting the wet zone. The horizontal line within each box represents the median. The lower and upper bounds of the box indicate the first and third quartiles, respectively. The lines (“whiskers”) extending from the boxes correspond to the minimum and maximum values. Outliers were identified as dots. Asterisks above paired box plots indicate *P* < 0.05; n.s. indicates no significant difference, Wilcoxon signed-rank exact tests.

Bed bugs spent substantially less time on the wet zone when compared to the dry zone (Wilcoxon signed-rank exact test, low: *V* = 371, *P* < 0.0001, medium: *V* = 378, *P* < 0.0001, high: *V* = 378, *P* < 0.0001, control *V* = 254, *P* = 0.12). The median cumulative durations spent in the wet and dry zones were 9.7 (IQR: 0-86.2) *vs*. 583.6 (IQR: 503.8-599.6) s (low), 0 (IQR: 0-0.5) *vs*. 600.0 (IQR: 596.6-600.0) s (medium), 0 (IQR: 0-3.5) *vs*. 600.0 (IQR: 595.5-600.0) s (high), 320.7 (IQR: 281.0-398.3) *vs*. 262.8 (IQR: 194.2-317.0) s (control), respectively (Figure 2B).

The mean speeds of the bed bugs were different between the wet and dry zones. For the lowest application rate, bed bugs walked significantly faster in wet than dry zones (Wilcoxon signed-rank exact test, *V* = 30, *P* = 0.014); the median of the mean speeds on the wet and dry zones were 0.32 (IQR: 0.24-0.45) *vs*. 0.24 (IQR: 0.19-0.36) cm/s, respectively. For the medium application rate, there was no difference in walking speed between wet and dry zones (Wilcoxon signed-rank exact test, *V* = 18, *P* = 1.0); the median of the mean speeds on the wet and dry zones were 0.23 (IQR: 0.08-0.35) *vs*. 0.17 (IQR: 0.14-0.35) cm/s, respectively. At the highest application rate, bed bugs walked significantly slower in wet than dry zones (Wilcoxon signed-rank exact test, *V* = 26, *P* = 0.047); the median of the mean speeds in the wet and dry zones were 0.26 (IQR: 0.16-0.30) *vs*. 0.36 (IQR: 0.28-0.37) cm/s. In the control, there was no difference in walking speed between the two dry zones (Wilcoxon signed-rank exact test, *V* = 168, *P* = 0.86); the median of the mean speeds on the two dry zones was 0.39 (IQR: 0.18-0.48) *vs*. 0.34 (IQR: 0.15-0.45) cm/s. Significant differences in walking speed for the low application rate were primarily driven by seven of eight adult females and six of eight adult males that walked faster on wet than dry zones and for the high application rate by six of seven bugs (data pooled) that walked slower on wet than dry zones (Supplemental Figure 1).

### Bed bugs increase speed after turning away from the wet zone

Turning behavior was observed in 436 of 610 approaches to the wet zone without touching it. Flee speed after turning away (*v*_*flee*_) was significantly greater than approach speed (*v*_*approach*_) at all three water application rates (Wilcoxon signed-rank tests with continuity correction): *V* = 2638, *P* < 0.0001 (low), *V* = 2875, *P* < 0.0001 (medium), *V* = 3498, *P* < 0.0001 (high) (Figure 3). The median of the respective approach and flee speeds were 0.34 (IQR: 0.27-0.51) *vs*. 0.44 (IQR: 0.29-0.78) cm/s (low), 0.38 (IQR: 0.28-0.55) *vs*. 0.43 (IQR: 0.31-0.63) cm/s (medium), and 0.34 (IQR: 0.26-0.45) *vs*. 0.43 (IQR: 0.29-0.64) cm/s (high). For all application rates, the speed difference (*v*_*flee*_ -*v*_*approach*_) differed significantly from zero (Wilcoxon signed rank test with continuity correction), mean ± SE: 0.13 ± 0.024 cm/s, *V* = 7373, *P* < 0.0001 (low), 0.10 ± 0.023 cm/s, *V* = 6716, *P* < 0.0001 (medium), 0.12 ± 0.019 cm/s, *V* = 11727, *P* < 0.0001 (high), and there was no difference among application rates (Kruskal-Wallis test, *χ*^*2*^= 1.51, *df* = 2, *P* = 0.47).

**Fig. 3.**
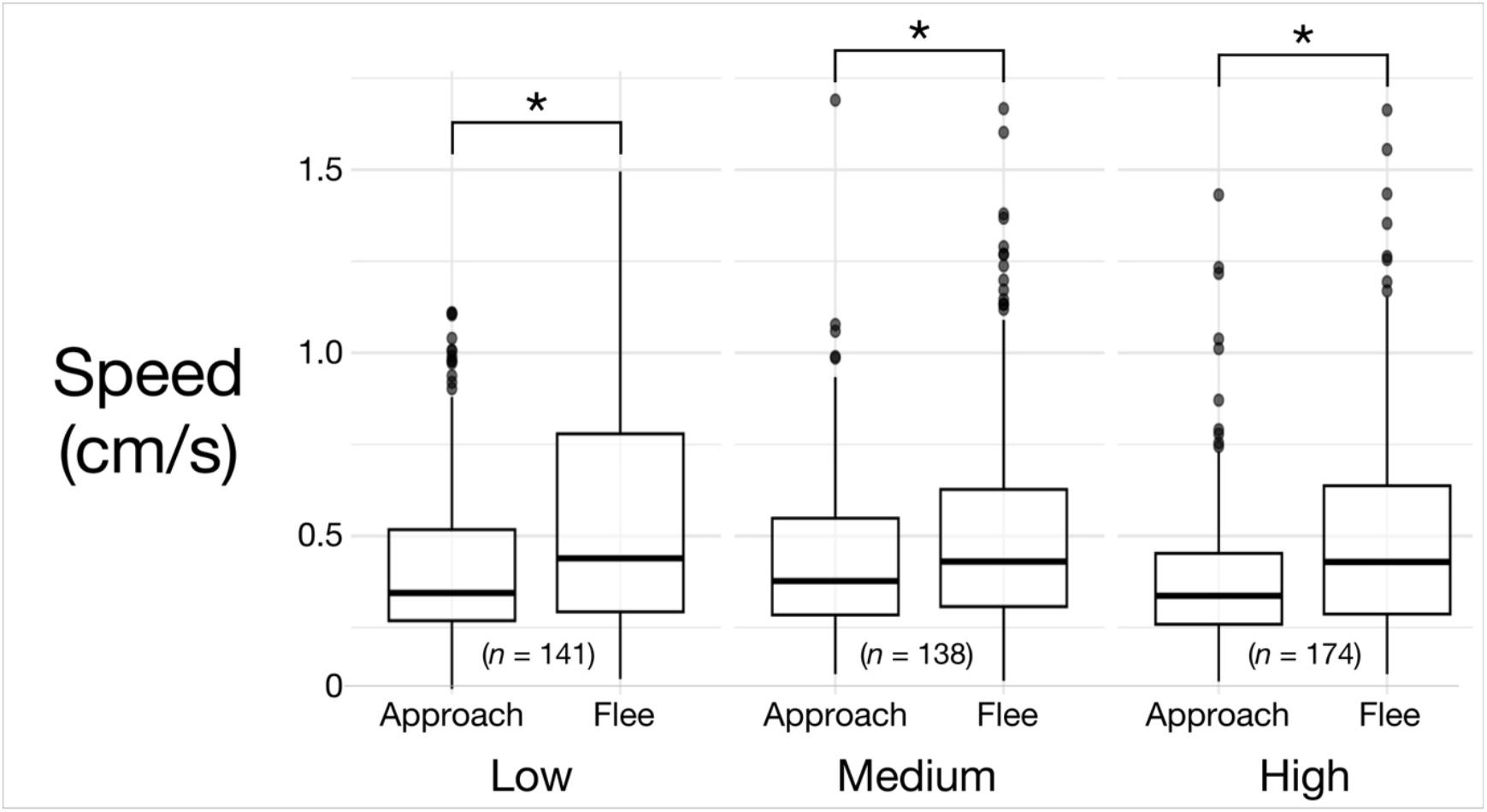
Box plots of approach speed (*v*_*approach*_) and flee speed (*v*_*flee*_) for all bed bugs. Flee speed after turning away from the wet zone was significantly greater than approach speed at all three water application rates (Wilcoxon signed-rank tests with continuity correction): *V* = 2638, *P* < 0.0001 (low); *V* = 2875, *P* < 0.0001 (medium); *V* = 3498, *P* < 0.0001 (high). The horizontal line within each box represents the median. The lower and upper bounds of the box indicate the first and third quartiles, respectively. The lines (“whiskers”) extending from the boxes correspond to the minimum and maximum values. Outliers are shown as individual dots. Sample sizes (number of turns without contacting the wet zone) are provided for each application rate: low (*n* = 141), medium (*n* = 138), and high (*n* = 174).

### Minimum distance to the wet zone before turning depends on water application rate and developmental stage

The minimum distance to the wet zone prior to turning for all application rates ranged from 5.39×10^−5^ cm to 1.55 cm. The minimum distance to the wet zone prior to turning for the lowest application rate was significantly shorter (Kruskal-Wallis test followed by Dunn’s test for multiple comparisons) than for the two higher rates (Figure 4A, *χ*^*2*^ = 9.86, *df* = 2, *P* = 0.0072). The median minimum distances before turning were 0.40 (IQR: 0.31-0.66) cm (low), 0.51 (IQR: 0.33-0.88) cm (medium), and 0.51 (IQR: 0.35-0.82) cm (high).

**Fig. 4.**
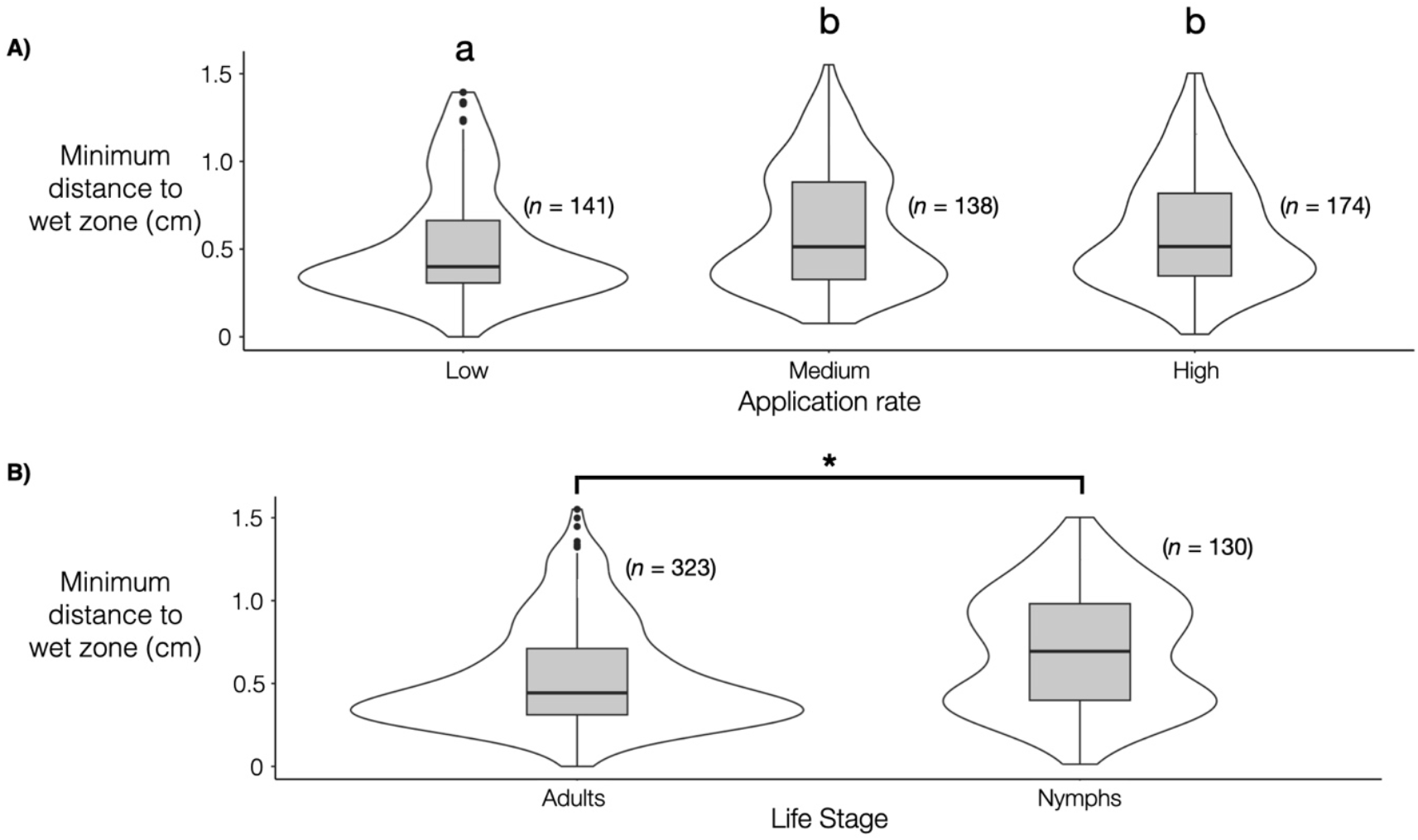
Minimum distance from the wet zone before making a turn by water application rate for all life stages pooled (A) and by life stage for all doses pooled (B). Box plots with different letters above (A) are significantly different (Kruskal-Wallis test followed by Dunn’s test for multiple comparisons with a Bonferroni correction, *χ*^*2*^ = 9.87, *df* = 2, *P* = 0.0072), with Dunn’s test for multiple comparisons (Bonferroni correction). Asterisk (B) indicates a significant difference between adults and nymphs (Wilcoxon rank sum test: *W* = 15170, *P* < 0.0001). The horizontal line within each box represents the median. The lower and upper bounds of the box indicate the first and third quartiles, respectively. The lines (“whiskers”) extending from the boxes correspond to the minimum and maximum values. Outliers were identified as dots. A violin plot surrounds each box plot to visually display the data distribution within each box plot. Sample size (*n*) denotes the number of turns bed bugs made without contacting the wet zone.

Nymphs made turns further away from the wet zone than adults (Figure 4B) (Wilcoxon rank-sum test, *W* = 15170, *P* < 0.0001; *n* = 323 and 130 for adults and nymphs, respectively). The median of the minimum distances to the wet zone before making a turn was 0.44 (IQR: 0.31-0.71) and 0.69 (IQR: 0.40-0.98) cm for adults and nymphs, respectively.

### Analysis of turning behavior over repeated approaches

To assess whether bed bugs change their turning behavior over repeated approaches to the wet zone, we calculated the slope of the minimum distance to the wet zone for each bed bug with two or more approaches. A negative slope would indicate that bed bugs turned closer to the wet zone over time, while a positive slope would indicate that they turned farther away. For all application rates, the slopes were not significantly different from zero (mean ± SE): -0.003 ± 0.24 cm/approach number (low), 0.055 ± 0.19 cm/approach number (medium), and 0.051 ± 0.26 cm/approach number (high) (Wilcoxon signed-rank exact tests, *P* > 0.05). This suggests that bed bugs did not systematically change their turning behavior over repeated approaches to the wet zone. There is no significant difference between application rates (Kruskal Wallis rank sum test, Kruskal-Wallis *χ*^*2*^ = 2.95, *df* = 2, *P* = 0.229).

## Discussion

This study demonstrated that common bed bugs avoid surfaces wet with deionized water. Remarkably, there was an 86.9% avoidance rate, with only 69 of 527 approaches resulting in direct contact with the wet zone. The observed behavior aligns with negative hydrotaxis, a well-documented orientation mechanism in arthropods, and likely serves as an adaptive strategy to mitigate environmental hazards such as fungal infection or drowning.

Importantly, our experimental design did not assess bed bug responses to relative humidity or orientation to water sources, which involve distinct sensory inputs and behavioral strategies. While antennal hygroreceptors mediate responses to ambient humidity (Steinbrecht and Müller 1976; Altner and Loftus 1985), our findings demonstrate that avoidance of wet surfaces integrates both antennal detection and tactile feedback from direct contact. This distinction is critical, as the sensory mechanisms underlying substrate wetness responses may differ from those involved in distant humidity detection or water-source orientation.

The variation in bed bug walking speeds across wet surfaces—higher speeds at low water application rates and lower speeds at high rates (Figure 2C) demonstrates context-dependent locomotion modulation akin to other arthropods (Ritzmann and Büschges 2007). Similar substrate-driven speed adjustments occur when cockroaches cross elastic surfaces (Spence et al. 2010) and stick insects navigate gaps (Blaesing and Cruse 2004). The faster speeds for bed bugs walking on lightly wet surfaces could indicate escape behavior to a stimulus similar to roaches responding to puffs of air (Ritzmann 1984). However, the inconsistent response in speed across wetness levels—including unchanged medium-rate speeds and slower high-rate speeds—shows this response is not purely escape-driven.

The faster fleeing speed after turning away from the wet zone (Figure 3) suggests that bed bugs perceive wet surfaces as a threatening stimulus, consistent with the ‘fast start’ fleeing behavior observed in many animals (Hale 1999; Wakeling et al. 1999; Hale et al. 2002). This behavior aligns with startle responses in other insects, such as acoustic startle responses in flying insects (Hoy et al. 1989) or looming stimuli in *Drosophila* (Card and Dickinson 2008). The lack of significant differences in speed difference (*v*_*flee*_ *-v*_*approach*_) across water application rates suggests that the degree of surface wetness does not influence the intensity of this avoidance behavior.

The distance at which bed bugs turned before contacting the wet zone (Figure 4A) indicates a sensitivity to moisture levels, with bed bugs turning closer to the wet zone at lower application rates. This pattern suggests that bed bug hygroreception may vary in sensitivity depending on the moisture levels in the surrounding air, consistent with the idea that insects modulate their responses based on stimulus intensity (Taylor and Krapp 2007). Future research should explore how bed bugs integrate these sensory inputs when responding to different water-related stimuli. Comparative studies of their behavior toward high humidity versus direct water contact, or their ability to detect water sources at a distance, would provide a more complete understanding of their hydrotactic responses and potentially reveal new opportunities for pest management.

The finding that nymphs turned farther away from the wet zone than adults (Figure 4B) suggests developmental differences in hygroreceptor sensitivity, potentially linked to differences in cuticle sclerotization and vulnerability to fungal pathogens (Hepburn and Roberts 1975; Hepburn and Chandler 1976; Vincent and Wegst 2004; Li et al. 2020). While these findings provide an initial basis for understanding hygroreceptor sensitivity in bed bugs, further research is required to explore this potential developmental difference. While the location of hygroreceptors in arthropods varies, including the antennae, legs, mouthparts, and wings (Gunn 1937; Altner and Loftus 1985; Gaffin et al. 1992; Ehn and Tichy 1994; Kröber and Guerin 2000), it remains unclear at what point bed bugs acquire hygroreception in their development. Changes in chemoreceptor sensitivity over the course of insect development (Blaney et al. 1986) suggest that hygroreception might also vary throughout life stages. These results provide an initial basis for understanding developmental differences in hygroreceptor sensitivity, and further research is needed to explore this potential variation.

The analysis of turning behavior over repeated approaches to the wet zone revealed no systematic change in the minimum distance at which bed bugs turned away. This suggests that bed bugs exhibit a consistent avoidance response to wet surfaces, regardless of prior encounters. Such behavioral consistency may reflect the evolutionary pressures of their natural and human-associated habitats, where wet surfaces pose a consistent threat (e.g., fungal infection, drowning). In contrast, tick larvae (*B. microplus*) in laboratory settings eventually accept wet surfaces after repeated approaches (Kröber and Guerin 2000). The differing responses of bed bugs and ticks highlight how evolutionary history and ecological context shape behavioral strategies in arthropods. Further research could explore whether bed bugs exhibit similar behavioral consistency in response to other environmental stimuli, such as heat or chemical cues.

Avoiding wet surfaces likely reduces the risk of fungal infections for bed bugs, a behavior that may be a conserved trait across Cimicidae, given its prevalence in multiple species. This ecological strategy is supported by both the vulnerability of bed bugs to fungal entomopathogens in high-humidity environments (Pasanen et al. 1991; Ashbrook et al. 2022) and analogous susceptibility patterns in other hematophagous insects, such as *Rhodnius prolixus* eggs, under 100% relative humidity (Schilman 1998). The evolutionary roots of this behavior are further underscored by the ecology of ancestral bed bugs, which inhabited caves and aggregated away from wet surfaces while feeding on bats (Usinger 1966). These findings highlight moisture avoidance as a critical survival strategy with deep phylogenetic significance in bed bug ecology, and further work could clarify whether modern human-associated lineages (e.g., *C. lectularius* vs. *C. hemipterus*) retain this trait to the same degree as their cave-dwelling ancestors.

The findings of this study have important implications for bed bug management, particularly with liquid-based chemical pesticides. Overapplying these pesticides may lead bed bugs to temporarily or persistently avoid treated areas, especially in high-humidity environments where surfaces remain wet longer (Rohwer 1931). This avoidance behavior could limit the areas available for bed bug movement and influence their future aggregation and harborage sites. The recommended application rate for products like CrossFire® Bed Bug Concentrate (1 gallon/1,000 square feet or 0.004 mL/cm^2^) is half the lowest rate used in this study (0.008 mL/cm^2^), where bed bugs predominantly avoided wet surfaces. Following these guidelines can help minimize the dispersal of bed bugs to untreated areas due to moisture in the spray.

## Acknowledgments

We thank Kathleen Campbell for maintaining the laboratory colony of bed bugs and providing feedback on the penultimate draft of the manuscript.

## Author contributions

Both authors contributed to the study’s conception and design. Material preparation, data collection, and analysis were performed by J.B.J. The first draft of the manuscript was written by J.B.J., and both authors commented on all versions of the manuscript. Both authors read and approved the final manuscript.

## Statements and Declarations

The authors declare no competing interests nor conflicts of interest.

## Supplemental Figures

**Supplemental Fig. 1.**
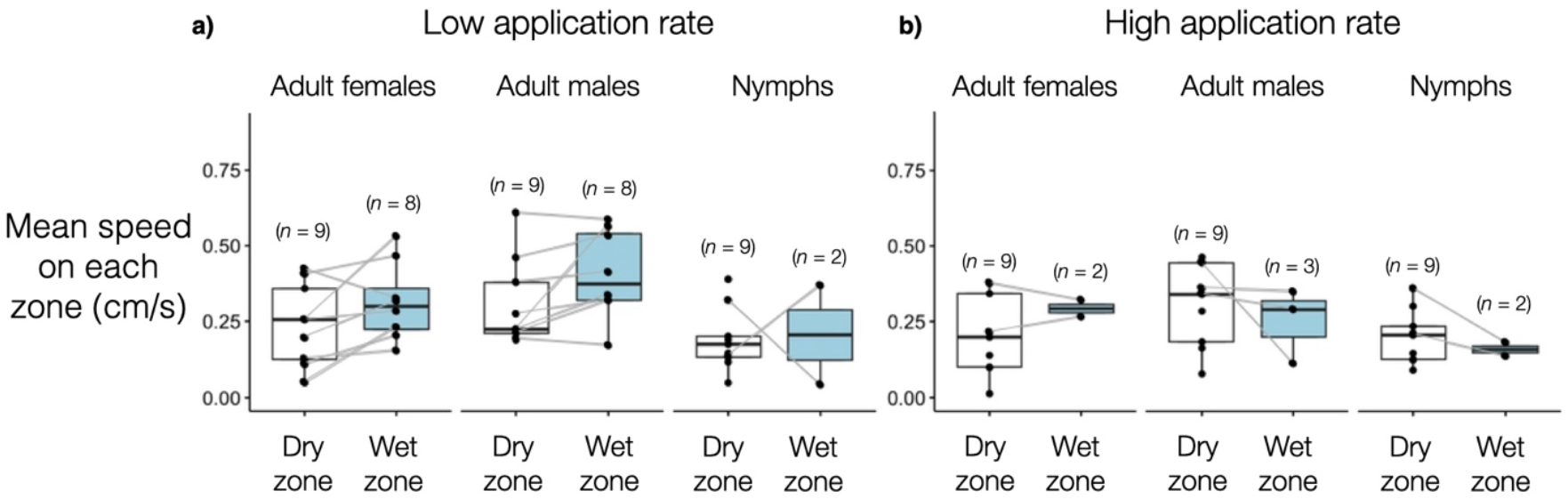
The mean speed of bed bugs on each zone (in cm/s) for the low and high application rates. For the low application rate, 7 of 8 adult females and 6 of 8 adult males walked faster on the wet zone relative to their speed on the dry zone. For the high application rate, 6 of 7 bed bugs walked slower on the wet zone relative to their speed on the dry zone. Sample size denotes the number of bed bugs.

**Supplemental Fig. 2.**
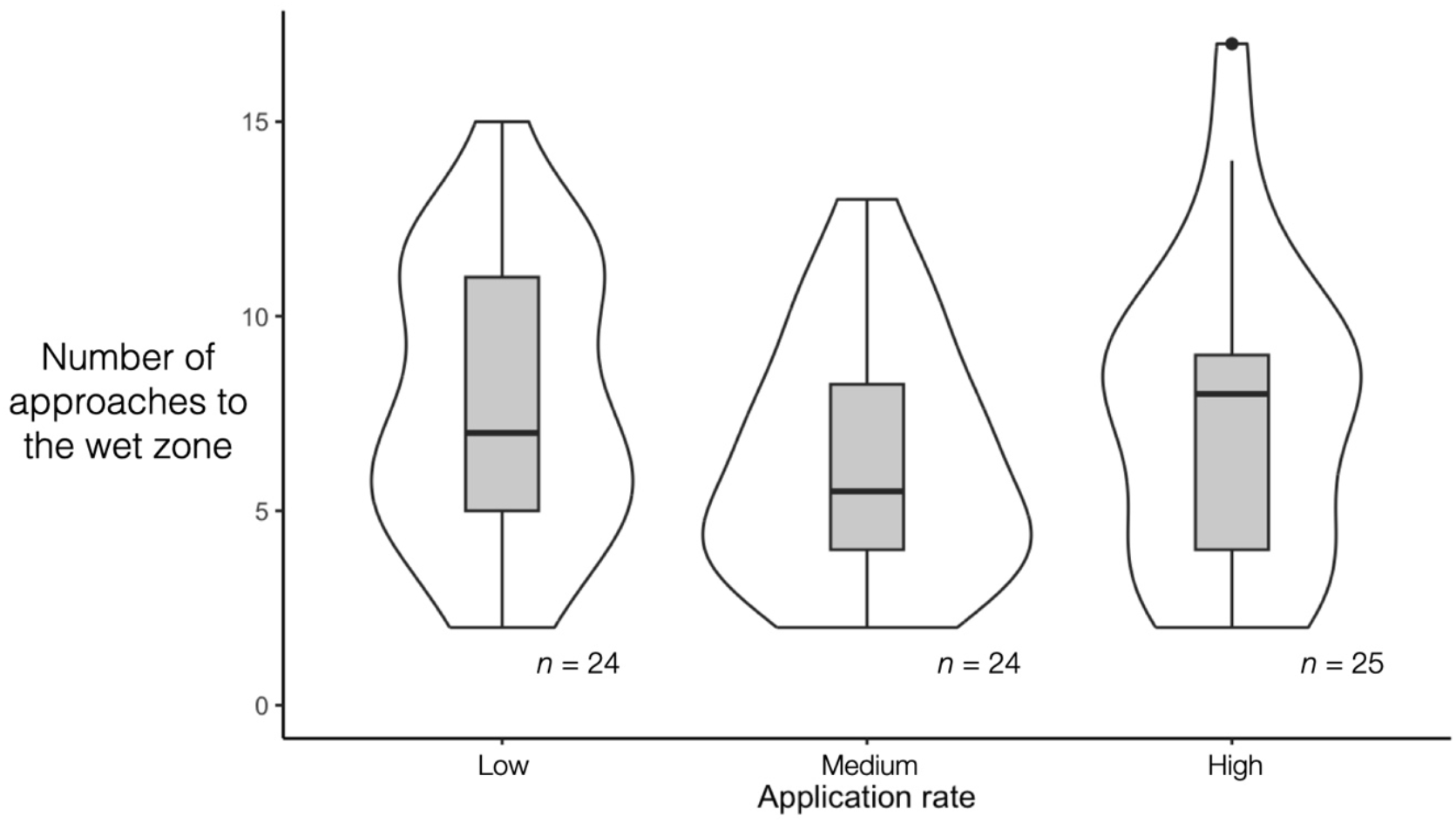
The number of approaches of bed bugs to the wet zone for all application rates. These approaches include instances where bed bugs contacted the wet zone. There is no significant difference between application rates (Kruskal Wallis rank sum test, Kruskal-Wallis *χ*^*2*^ = 1.96, *df* = 2, *P* = 0.38). The horizontal line within each box represents the median. The lower and upper bounds of the box indicate the first and third quartiles, respectively. The lines (“whiskers”) extending from the boxes correspond to the minimum and maximum values. Outliers were identified as dots. A violin plot surrounds each box plot to display the data distribution within each box plot visually. Sample size denotes the number of bed bugs.

